# Host transcriptomic profiling of COVID-19 patients with mild, moderate, and severe clinical outcomes

**DOI:** 10.1101/2020.09.28.316604

**Authors:** Ruchi Jain, Sathishkumar Ramaswamy, Divinlal Harilal, Mohammed Uddin, Tom Loney, Norbert Nowotny, Hanan Alsuwaidi, Rupa Varghese, Zulfa Deesi, Abdulmajeed Alkhajeh, Hamda Khansaheb, Alawi Alsheikh-Ali, Ahmad Abou Tayoun

## Abstract

Characterizing key molecular and cellular pathways involved in COVID-19 is essential for disease prognosis and management. We perform shotgun transcriptome sequencing of human RNA obtained from nasopharyngeal swabs of patients with COVID-19, and identify a molecular signature associated with disease severity. Specifically, we identify globally dysregulated immune related pathways, such as cytokine-cytokine receptor signaling, complement and coagulation cascades, JAK-STAT, and TGF-β signaling pathways in all, though to a higher extent in patients with severe symptoms. The excessive release of cytokines and chemokines such as *CCL2, CCL22, CXCL9* and *CXCL12* and certain interferons and interleukins related genes like *IFIH1, IFI44, IFIT1* and *IL10* were significantly higher in patients with severe clinical presentation compared to mild and moderate presentations. Moreover, early induction of the TGF-β signaling pathway might be the primary cause of pulmonary fibrosis in patients with severe disease. Differential gene expression analysis identified a small set of regulatory genes that might act as strong predictors of patient outcome. Our data suggest that rapid transcriptome analysis of nasopharyngeal swabs can be a powerful approach to quantify host molecular response and may provide valuable insights into COVID-19 pathophysiology.

## Introduction

Since the first case reported in December 2019 in Wuhan, SARS-CoV-2 has spread very quickly across 188 different countries leading to over 31,887,485 COVID-19 cases and 976,789 associated deaths worldwide as of 24^th^ September 2020.^1^ The common signs and symptoms of SARS-CoV-2 include fever, cough, fatigue, shortness of breath, and chest CT abnormalities.^2^ Some patients with COVID-19 quickly develop severe pulmonary symptoms including acute respiratory distress syndrome (ARDS), pulmonary edema, intense kidney injury and multiple organ failure^3^ whereas other patients present with no symptoms or with only mild disease.^4^ Pathogen load and prevalence of infections are higher in males than females demonstrating that sex physiology plays a role in infectious disease pathogenesis.^5^ Meng and colleagues^6^ showed that men who died of COVID-19 had elevated levels of systemic inflammatory markers such as neutrophil-to-lymphocyte ratio and C-reactive protein. Nevertheless, the mechanism behind increased mortality among older adults and males with COVID-19 remains speculative.

Despite the worldwide spreading, the host immune response against SARS-CoV-2 infection remains poorly characterized. Dysregulation of host immune response and activation of inflammatory cytokines, known as the “cytokine storm”, is associated with disease severity and poor prognosis.^7, 8, 9^ A recent host transcriptome study on patients with COVID-19 revealed distinct host inflammatory cytokine profiles and highlighted the association between SARS-CoV-2 pathogenesis and excessive release of cytokine such as CCL2/MCP-1, CXCL10/IP-10, CCL3/MIP-1A, and CCL4/MIP1B.^10^ Additionally, Lieberman *et al*. showed upregulation of antiviral factors such as OAS1-3 and IFIT1-3, and Th1 chemokines CXCL9/10/11 upon SARS-CoV2 induced antiviral response. This study also showed immune responses may underlie disparities between males and females, and among the elderly compared to younger age groups.^11^

At present, it is not known if the host gene expression profile varies among patients with mild, moderate, or severe clinical outcomes. Identification of such transcriptomic differences can be useful for predicting COVID-19 outcomes and for better management and earlier interventions especially for patient groups with severe outcomes. In this study, we performed RNA sequencing on nasopharyngeal tissue of patients with mild, moderate, and severe disease to characterize transcriptomic regulation and immune response differences among these patients, and to identify markers for early identification of patients with possible severe disease outcomes.

## Methods

### Patient cohort and ethics statement

This study was approved by the Dubai Scientific Research Ethics Committee -Dubai Health Authority (approval number #DSREC-04/2020_02). The Ethics committee waived the requirement for informed consent since this study was part of a public health surveillance and outbreak investigation in the UAE. The electronic medical records of patients with laboratory confirmed SARS-CoV-2 were reviewed and important clinical data were extracted using the World Health Organization (WHO) case report form.

### Sample preparation, RNA isolation, library construction and sequencing

RNA was extracted from nasopharyngeal swabs using the QIAamp Viral RNA Mini or the 213 EZ1 DSP Virus Kits (Qiagen, Hilden, Germany). All patients in this study tested positive for SARS-CoV-2 by RT-qPCR performed at Dubai Health Authority Hospitals. RNA libraries were then prepared for shotgun transcriptome sequencing using the TruSeq Stranded Total RNA Library kit from Illumina (San Diego, CA, USA) as previously described.^12,13,14^ Libraries were then sequenced using the NovaSeq SP Reagent kit (2 × 150 cycles) from Illumina (San Diego, CA, USA) to generate a minimum of 15M reads per sample (**Supplementary Table 1**).

**Table 1.** Clinical Characteristics of COVID-19 patients in this study.

| Characteristic | All Patients (N=50) | Stratified by Disease Severity |  |  |
| --- | --- | --- | --- | --- |
|  |  | Asymptomatic/Mild (N=37) | Moderate (N=10) | Severe/Critical (N=3) |
| Age Mean (SD), yr | 40.92 (16.03) | 36.4 (14.04) | 49.28 (14.35) | 68 (3.56) |
| Female Sex - no./total no. (%) | 18/50 (36%) | 16/37 (43.24%) | 1/10 (10%) | 1/3 (33.33%) |
| Body Mass (kg) - Mean (SD) | 66.2 (30.84) | 63.4 (28.65) | 73.07 (41.83) | 77.93 (8.29) |
| Current Smoker - no./total no. (%) | 0/50 (0%) | 0/37 (0%) | 0/10 (0%) | 0/3 (0%) |

| **Coexisting Disorder -**<br>**no./total no. (%)** |  |  |  |  |
| --- | --- | --- | --- | --- |
| Chronic Cardiac Disease (not Hypertension) | 3/50 (6%) | 1/37 (2.7%) | 2/10 (20%) | 0/3 (0%) |
| Hypertension | 9/50 (18%) | 5/37 (13.51%) | 3/10 (30%) | 1/3 (33.33%) |
| Chronic Pulmonary Disease | 0/50 (0%) | 0/37 (0%) | 0/10 (0%) | 0/3 (0%) |
| Asthma | 0/50 (0%) | 0/37 (0%) | 0/10 (0%) | 0/3 (0%) |
| Chronic kidney disease | 1/50 (2%) | 0/37 (0%) | 0/10 (0%) | 1/3 (33.33%) |
| Chronic liver disease | 0/50 (0%) | 0/37 (0%) | 0/10 (0%) | 0/3 (0%) |
| Chronic Neurological Disorder | 0/50 (0%) | 0/37 (0%) | 0/10 (0%) | 0/3 (0%) |
| Diabetes Mellitus | 8/50 (16%) | 3/37 (8.1%) | 3/10 (30%) | 2/3 (66.66%) |
| Malignant Neoplasms | 0/50 (0%) | 0/37 (0%) | 0/10 (0%) | 0/3 (0%) |

### Data analysis

The sequencing read quality was checked using FastQC v0.11.8^15^ and MultiQC v1.6.^16^ High quality reads (Q ≥ 30) were first mapped to rRNA sequences to remove potential rRNA reads using hisat v2-2.1.0^17^ with the default parameter. The remaining reads were then mapped to the GRCh37 (hg19). The unmapped reads were then mapped to the SARS-CoV-2 genome (GenBank Accession number: NC_045512.2) using BWA v0.7.17. PCR duplicates were removed with Picard MarkDuplicates program v2.18.17.^18^ Samples with more than 6M reads aligned to the GRCh37 (hg19) were considered for further analysis. RNA sequencing data obtained from nasopharyngeal swabs of samples confirmed to be COVID-19 negative by RT-qPCR, hereafter referred to as controls (n = 32), were downloaded from GSE152075.^11^

The number of reads mapped to each gene in the genome (GRCh37) was calculated using the FeatureCounts program in the SubReads package v2.0.1.^19^ DESeq2 package v1.28.1^20^ was applied to perform batch effects and normalization. In brief, DESeq2 uses the median of ratios method to normalize data and estimate size factors (which control for differences in the library size of the sequencing experiment). Further, gene expression change was calculated with respect to control for each groups using Wald test and *p-*values and log_2_ fold change was extracted. The resulting genes *p*-value was adjusted using the Benjamini and Hochberg method. These adjusted *p*-value (adj *p-*value) are calculated as: gene *p*-value × (m/i)) where *m* is the total number of genes, *i* is the gene *p*-value rank. Here we have considered a fraction of 5% false positives acceptable hence the genes with adj *p*-value < 0.05 were called as significant.

Pathway enrichment analysis was performed using the clusterProfiler package v 3.16.0^21^ to identify shared pathways among DEGs. Pathways with adj *p*-value < 0.05 were reported as significant. Heatmaps were generated using Morpheus^22^ and volcano plots were generated using VolcaNoseR package.^23^ Both case and control sequencing data were normalized to counts per million (CPM), which is calculated by normalizing the read counts for a given gene by the total counts per sample. CPM values were used to compare expression between certain genes and violin plots were generated using GraphPad Prism v8.0. ^24^ Two tailed Mann–Whitney U test *P*-values are reported. A schematic illustration of data analysis is represented in **Figure 1**.

**Figure 1.**
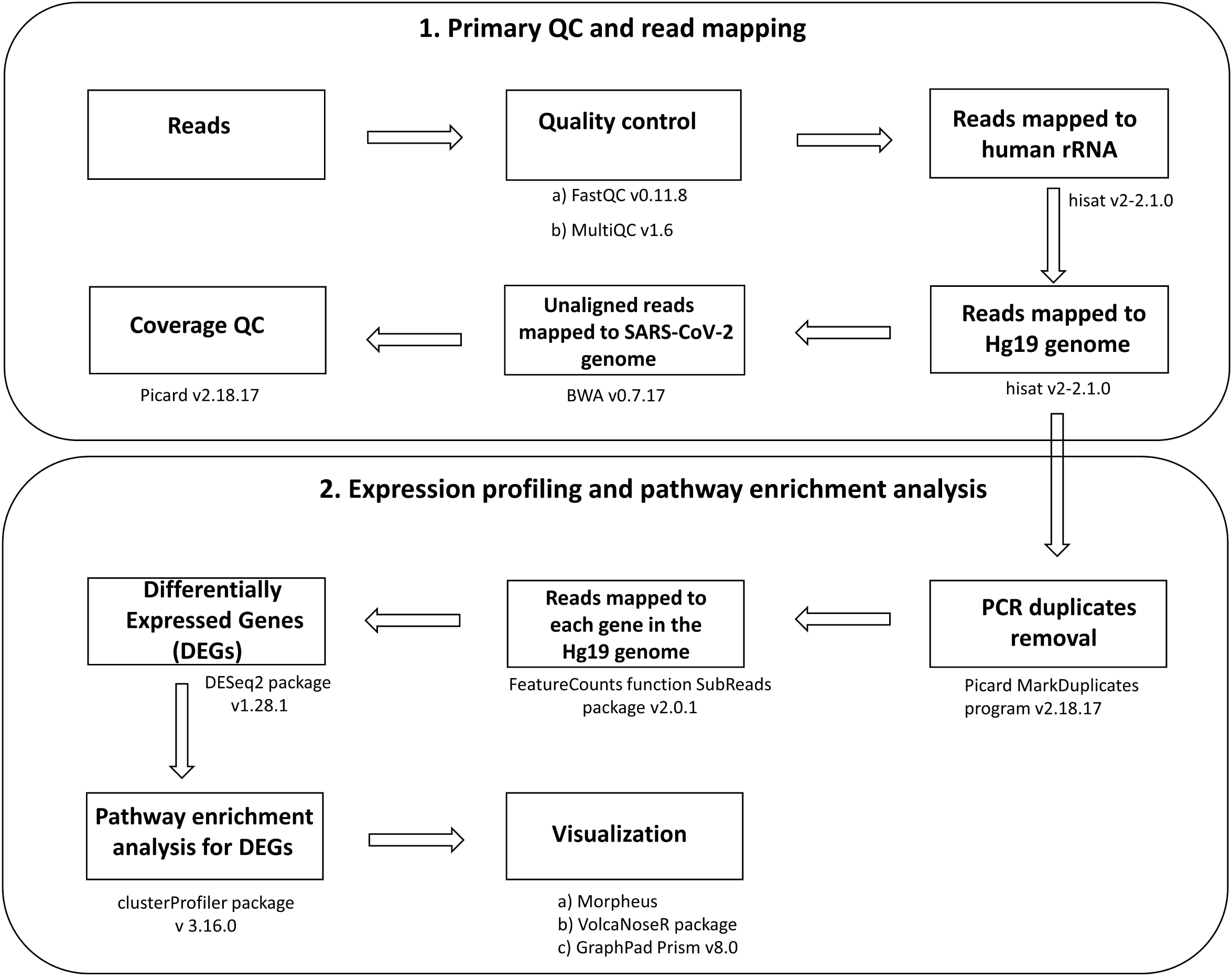
Schematic workflow for Transcriptomic analysis. Sequencing data underwent pre-processing which includes primary QC and read mapping, followed by differenticially expressed genes (DEG) analysis and downstream pathway enrichment analysis and visualization.

## Results

### Patient Cohort

Our patient cohort consisted of fifty patients with COVID-19 (36% female) who tested positive for SARS-CoV-2 by RT-qPCR with a mean age of 40.9 years (ranging from 3 – 70 years) at diagnosis. Cases were categorized into mild, moderate, and severe as previously described.^12^ Briefly, mild cases were asymptomatic or had mild non-life-threatening symptoms, while moderate cases presented with symptoms (such as persistent fever) requiring medical attention and/or hospitalization. Severe cases presented with advanced disease and pneumonia requiring admission to intensive care units and life-support treatment (such as mechanical ventilation). In total, 37 patients (21 males and 16 females; mean age 36.4 years) had mild disease, 10 patients (9 males and 1 female; mean age 49.3 years) had moderate presentations, while 3 patients (2 males and 1 female; mean age 68.0 years) had severe/critical disease. All patients were non-smokers. All 3 patients with severe disease had chronic health issues (hypertension, chronic kidney disease and/or diabetes mellitus) (**Table 1**).

### Global transcriptomic changes in patients with COVID-19

To understand the host mechanisms of SARS-CoV-2 infections, we explored transcriptomic profiling of patients with COVID-19 using RNA extracted from nasopharyngeal samples (see Methods). We first mapped high quality reads to the human genome and the SARS-CoV-2 genome (GenBank accession number: NC_045512.2) and summarized sequencing statistics in **Supplementary Table 1**. On average, ∼2% (ranging from 0.1% -13.6%) of the reads mapped to the SARS-CoV-2 genome confirming the presence of the virus, albeit coverage over the viral genome varied across patients most likely due to differences in viral loads. On the other hand, ∼65% (ranges from 24.3% -91.0%) of the RNA reads mapped to the human genome, enabling host transcriptomic analysis.

To detect global signature genes in patients with COVID-19 we combined transcriptomic data from patients with mild, moderate, and severe disease and made a comprehensive cohort called “COVID” to compare against controls for DEG analysis (see Methods). We detected 3572 upregulated genes and 445 downregulated genes with adj *p*-value□<□0.05 and fold-change□cutoff >□2. Several interferon, cytokine and immune-related genes, such as *CXCL5, CXCL12, CCL2, IFIH1, IFI44, IFIT1* and *IL10* were upregulated, whereas metabolic pathways or housekeeping genes, like *RPL41, RPL17, SLC25A6, CALM1* and *TUBA1A*, were downregulated (**Figure 2**).

**Figure 2.**
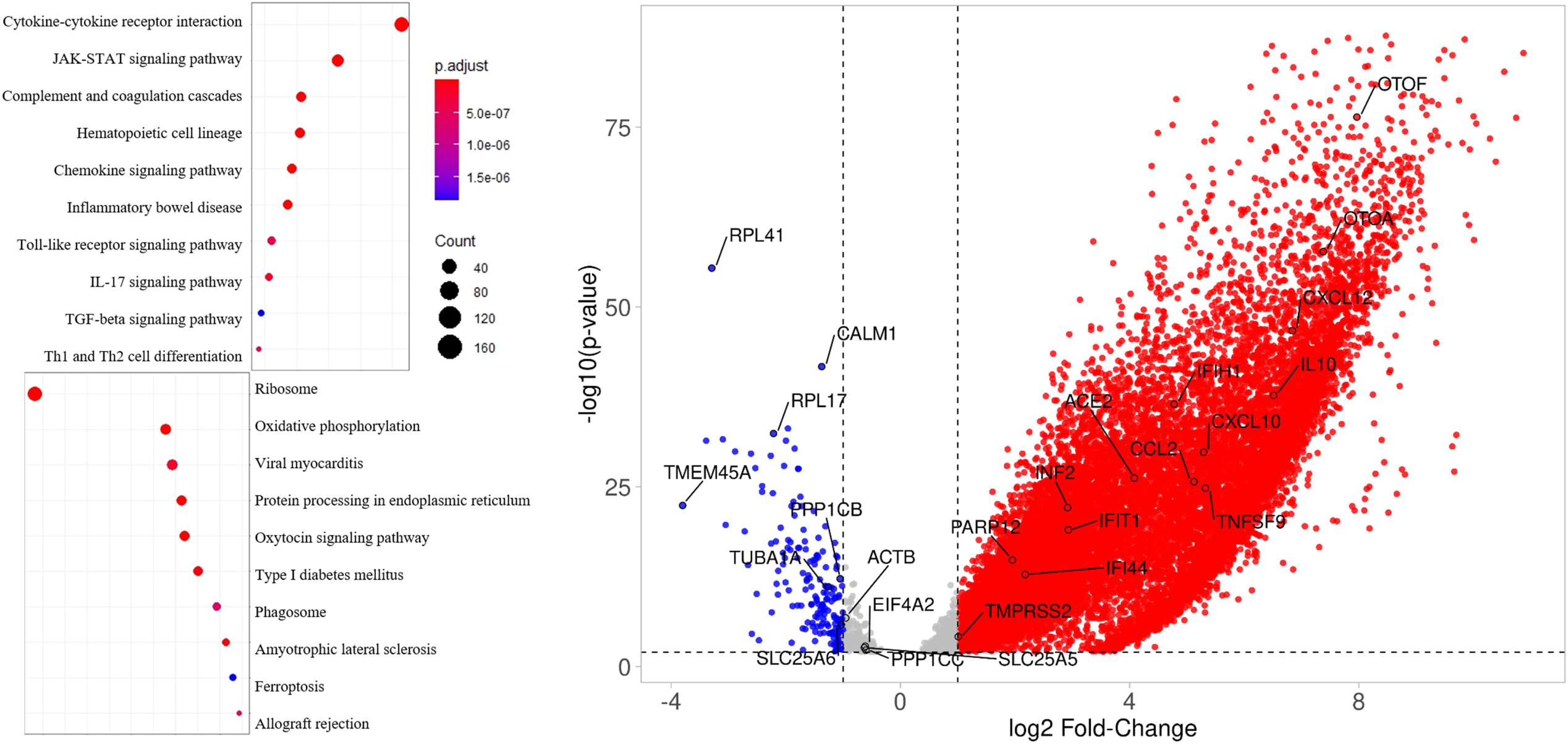
*Left*, Dotplot visualization of enriched Pathway terms in all COVID-19 patients. The color of the dots represents adj *p*-value for each enriched pathway, and size represents the percentage of genes enriched in the total gene set. *Right*, Volcano plot representing upregulated and downregulated genes. X-axis represents log_2_ fold change of genes and Y-axis represents –log_10_ *P*-value in differentially expressed gene (DEG) analysis.

Pathway enrichment analysis of the DEGs was performed to interrogate signaling programs induced by SARS-CoV-2. Up-regulated genes were related to cytokine-cytokine receptor interaction, JAK-STAT signaling pathway, complement and coagulation cascades and other inflammatory pathways. Whereas, several metabolic pathways were negatively enriched, including ribosome, oxidative phosphorylation and ER protein processing, which suggests a global reduction in the production of proteins related to cellular energy production (**Figure 2**).

### Transcriptome analysis of patients with mild, moderate and severe COVID-19

Next, we compared the transcriptomic profiles of patients with mild, moderate, and severe disease each separately with controls to determine if the above global signature, and the identified cytokine storm, hold similarly or varies by disease severity. Overall, we detected 2240 upregulated and 422 downregulated genes in mildly, 3447 upregulated and 302 downregulated genes in moderately, and 2691 upregulated and 215 downregulated genes in severely ill patient samples. Most of the immune response genes were significantly upregulated in all patient groups (**Supplementary Figure 1**).

Furthermore, pathway enrichment analysis showed modulation, to varying extents (**Figure 3**), of several overlapping pathways among mildly, moderately, and severely affected patients (**Supplementary Figure 1)**. In addition to upregulation of immune-related response genes, there was consistent disruption of the ribosome pathway. The SARS-CoV-2 receptor ACE2 is an interferon-regulated gene and is upregulated in response to SARS-CoV-2 infection.^25^ We also found that ACE2 expression was highest in patients with severe disease compared to patients with mild and moderate disease (**Figure 3**).

**Figure 3.**
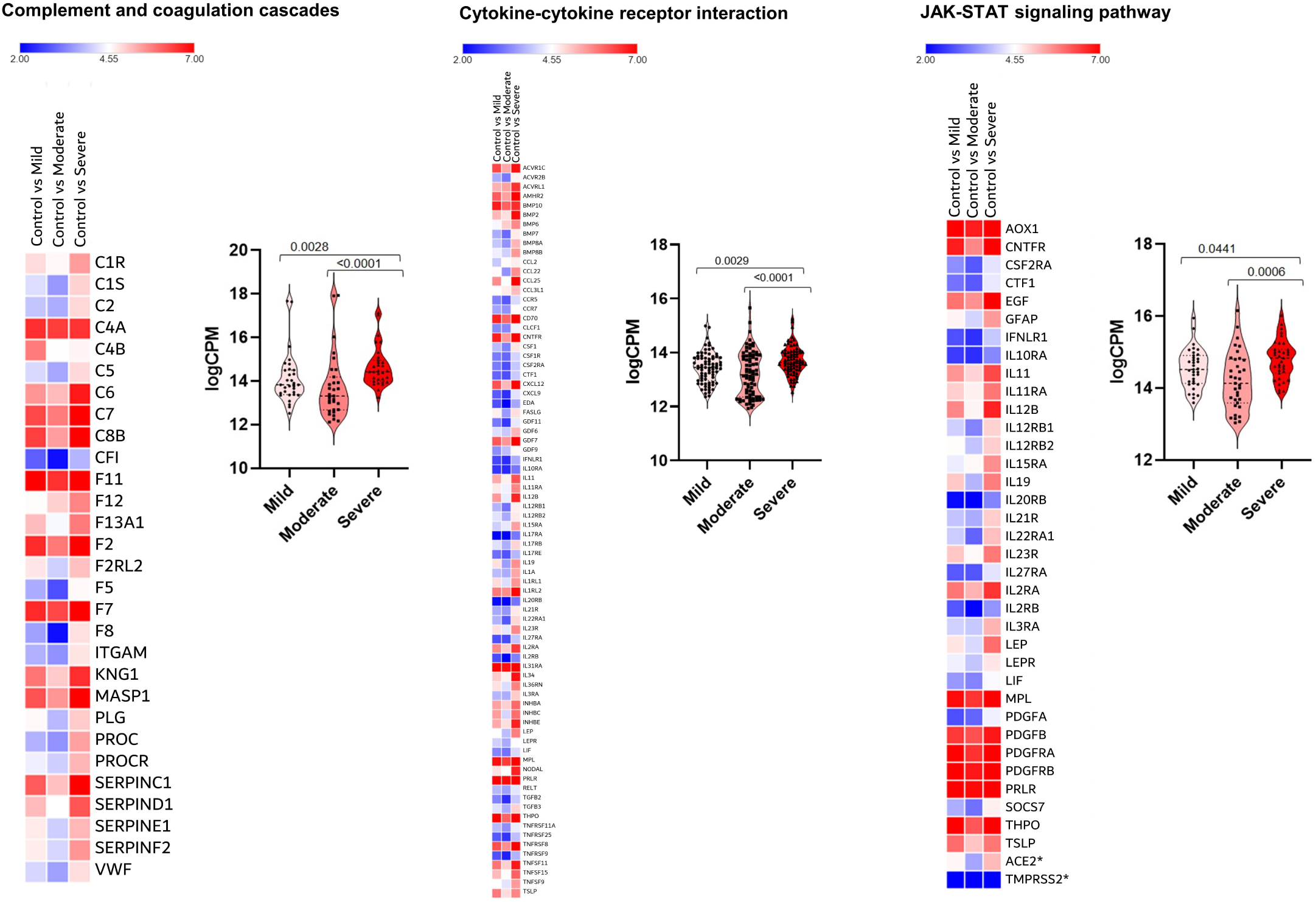
Heat map and Violin plots of pathways in patients with mild, moderate or severe disease. Pathways are: a) Complement and coagulation cascades, Cytokine-cytokine receptor interaction, JAK-STAT signaling pathway, d) TGF-beta signalling, Platelets activation, and Ribosome. Heat map depicts the log_2_ fold change of differentially expressed gene (DEGs) of COVID-19 patients compared with controls. Genes included have a log_2_ fold change of more than 1 and a *p*-adjusted value of less than 0.05. For every gene in a given pathway, the average counts per million (CPM) was calculated in cases with mild, moderate and severe disease and these CPM values were plotted (Violin plots) indicating median and quartiles as well as minima and maxima bounds. *Expression of *ACE2* and *TMPRSS2*.

### Comparison of functional enrichment among patients with mild, moderate, and severe COVID-19

To further delineate the differences among these groups, we compared the expression levels of genes and associated pathways (complement and coagulation cascades, cytokine-cytokine receptor interaction, JAK STAT signaling, TGF beta signaling pathway, platelets activation and ribosome pathway) between patients with mild, moderate, and severe disease. Gene expression in most regulated pathways was significantly higher (adj *p*-value < 0.05) in patients with severe disease compared to those with mild or moderate disease (**Figure 3**). For example, expression of certain interleukins (such as *IL11, IL12, IL19, IL34*), interleukin receptors (like *IL10RA, IL21R* and *IL11RA*), chemokines and tumour necrosis factor genes (such as *CXCL12, CXCL9, CCL25, CCL2, CCR5, CCR7, TNFSF9, TNFSF15, TNFRSF25* and *TNFRSF9*) were significantly higher in patients with severe disease compared to patients with mild or moderate disease (adj *p*-value < 0.05; **Figure 3**). Furthermore, complement and coagulation cascades (e.g. *C2, C5, C6, F12* and *F8*) were activated to a higher extent in patients with severe disease. Finally, patients with severe disease had higher expression of *STAT4, STAT5A* and *STAT5B* which are important components of JAK-STAT pathway and play a critical role in the fate of T helper cells^26^.

The highest fold change (> 2) in patients with severe disease was among the pro-inflammatory cytokines and chemokine receptors *CCR1* (CXCL8/CXCL6 receptor), *CCR6, CCR22, CCR25, IL3RA, IL11, IL19*, and *IL21RA*, the TGF-β signaling genes *BMP2, BMP7, PDGFA*, and *TNFSF11*, and the complement and platelet cascade genes *C6, C7, F2, F5, SERPINC1*, and *SERPIND1* (**Figure 4**). This distinction is likely due to viral infection resulting in release of inflammatory responses to a higher extent in patients with severe disease as compared to patients with mild or moderate disease.

**Figure 4:**
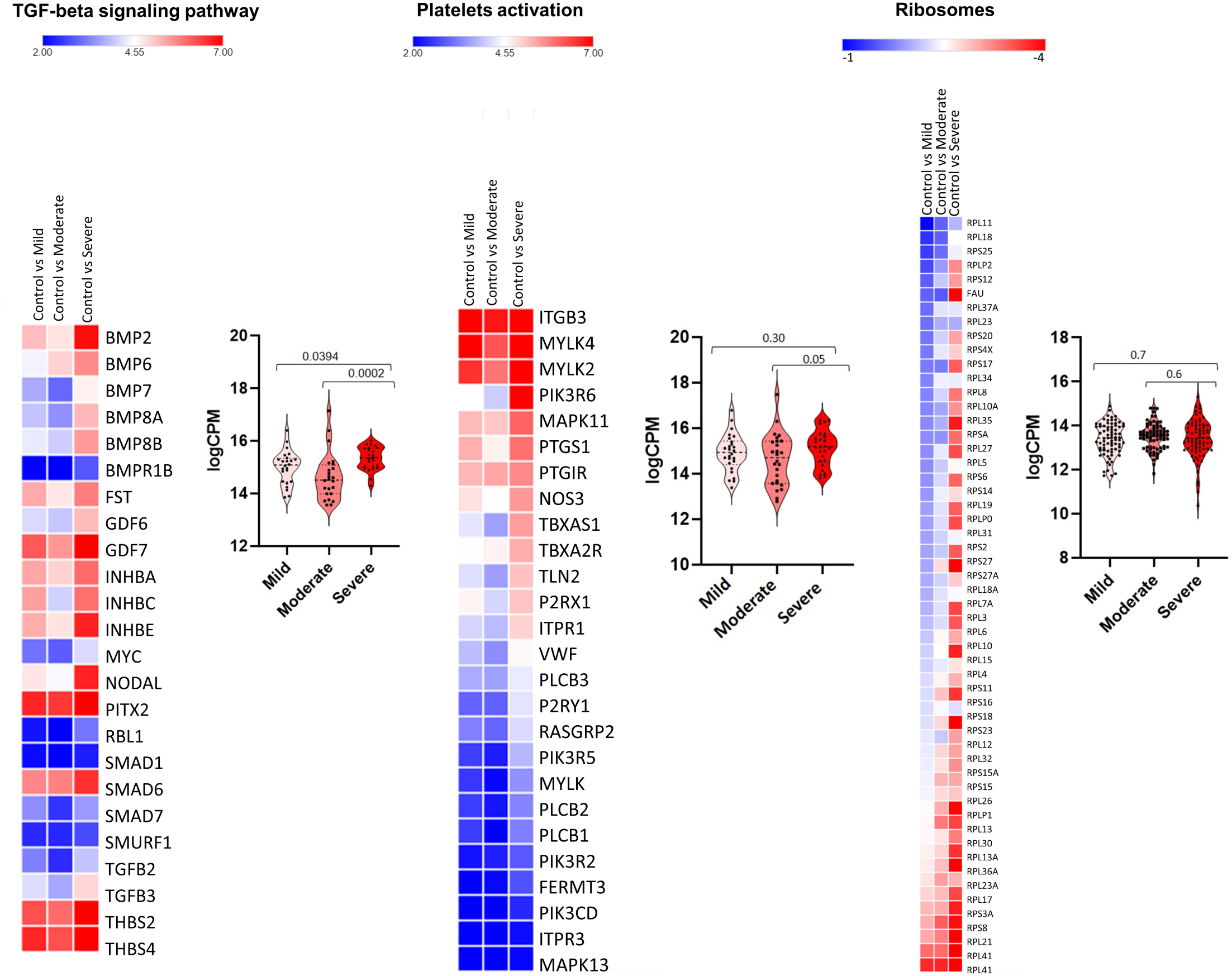
Scatter plot of fold change gene expression differences between patients with mild/moderate and severe outcomes. Fold change of mild/moderate was averaged and subtracted from severe fold change when compared to controls.

Finally, we compared transcriptomic changes in male and female COVID-19 patients with mild and moderate outcomes. Several immune related genes (*CSF2, TNFSF11, TNFRSF11B, IL18R1, IFIT1B* and *C4BPA*) were upregulated in male patients compared to females (**Figure 5**).

**Figure 5:**
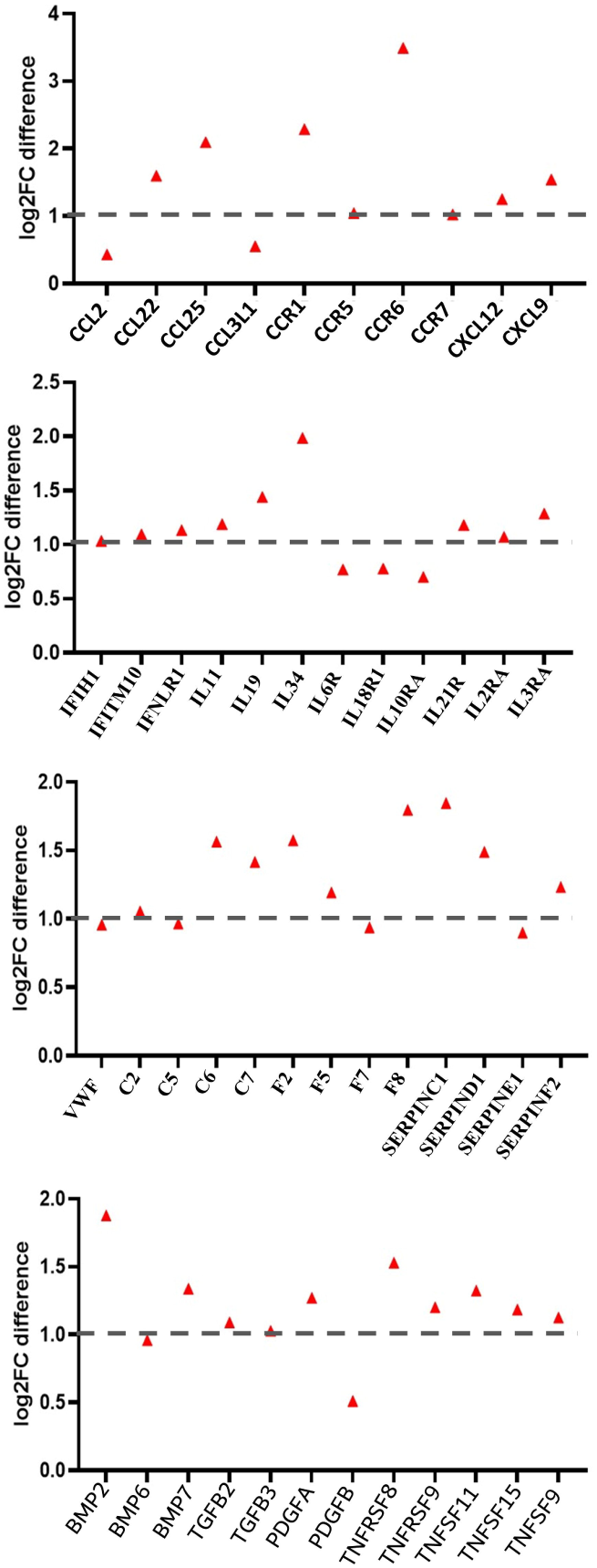
Heat map (top) and Violin plots (bottom) of differentially expressed genes in males and females. Heat map depicts the log_2_ fold change of differentially expressed gene (DEGs) of COVID-19 patients compared with controls. Genes included have a log_2_ (fold change) of more than 1 and adj *p*-value < 0.05. Violin plots represent median and 75th quartiles as well as minima and maxima bounds of CPM (counts per million) of highlighted differentially regulated genes.

## Discussion

Transcriptomic analysis of nasopharyngeal tissues from patients with COVID-19 reveals a robust early induction of cytokine and immune-related profiles by SARS-CoV-2 infection. In addition, elevated levels of complement and coagulation cascades were seen. Our findings are similar to those recently observed by Butler et al.^27^ in New York (United States) and Qin et al.^28^ in Wuhan (China). The latter work reported that an increase in ‘serum’ cytokine and chemokine levels, as well as in neutrophil-lymphocyte-ratio (NLR) in SARS-CoV-2 infected patients is correlated with the severity of the disease and adverse outcomes, suggesting a role for hyper-inflammatory responses in COVID-19 pathogenesis. Similarly, inflammation-induced coagulation pathways, which can themselves be regulated by the complement system, are pivotal in controlling pathogenesis associated with infections.^29^

Our RNA work, using nasopharyngeal tissue, extends those findings to show that those pathways are more significantly altered in patients with severe disease outcomes. Specifically, SARS-CoV-2 seems to induce expression of certain genes (**Figure 4**) to higher extents in patients with severe, compared to those with mild or moderate disease. This finding has direct prognostic and therapeutic implications. Identifying molecules targeting the pathways where those genes are involved might help ameliorate or avoid adverse clinical outcomes due to SARS-CoV-2. Moreover, this early expression signature can be used to predict the clinical course of disease, so that required early interventions can be implemented. However, larger datasets are needed to validate this expression signature and to show that it can reliably predict disease outcomes in the general population.

Certain molecular pathways require specific attention. In the majority of patients with COVID-19, respiratory failure is the primary cause of death and is often associated with uncontrolled inflammatory responses, edema, and lung fibrosis.^30^ This lung fibrosis is mainly triggered by transforming growth factor-beta (TGF-β).^10^ In our analysis, the TGF-β pathway was significantly upregulated in patients with severe disease suggesting they are more prone to respiratory failure. The PI3K/Akt and JAK/STAT signaling pathways are two other noteworthy pathways given that several cellular and immune responses, including host cell immune response to counteract viral infection ^31^, activation of interferon-stimulated genes (ISGs) ^32^, and activation of several regulatory and pro-inflammatory cytokines^33^ converge on those two pathways.

Our gender-specific transcriptomic analysis revealed a few genes which were upregulated in male patients compared to females (**Figure 5**). It is highly likely that many other key factors, which are involved in sex-specific responses to SARS-CoV-2, were not detected in our study. It is possible that our cohort was not large enough to perform appropriate comparisons on a large number of patients adequately stratified for age, sex, and disease severity.

Taken together, our study highlights key molecular and functional pathways involved in COVID-19 pathogenesis and characterizes a specific early expression signature associated with severe disease outcomes due to SARS-CoV-2.

## Figure Legends

**Supplementary figure 1:**
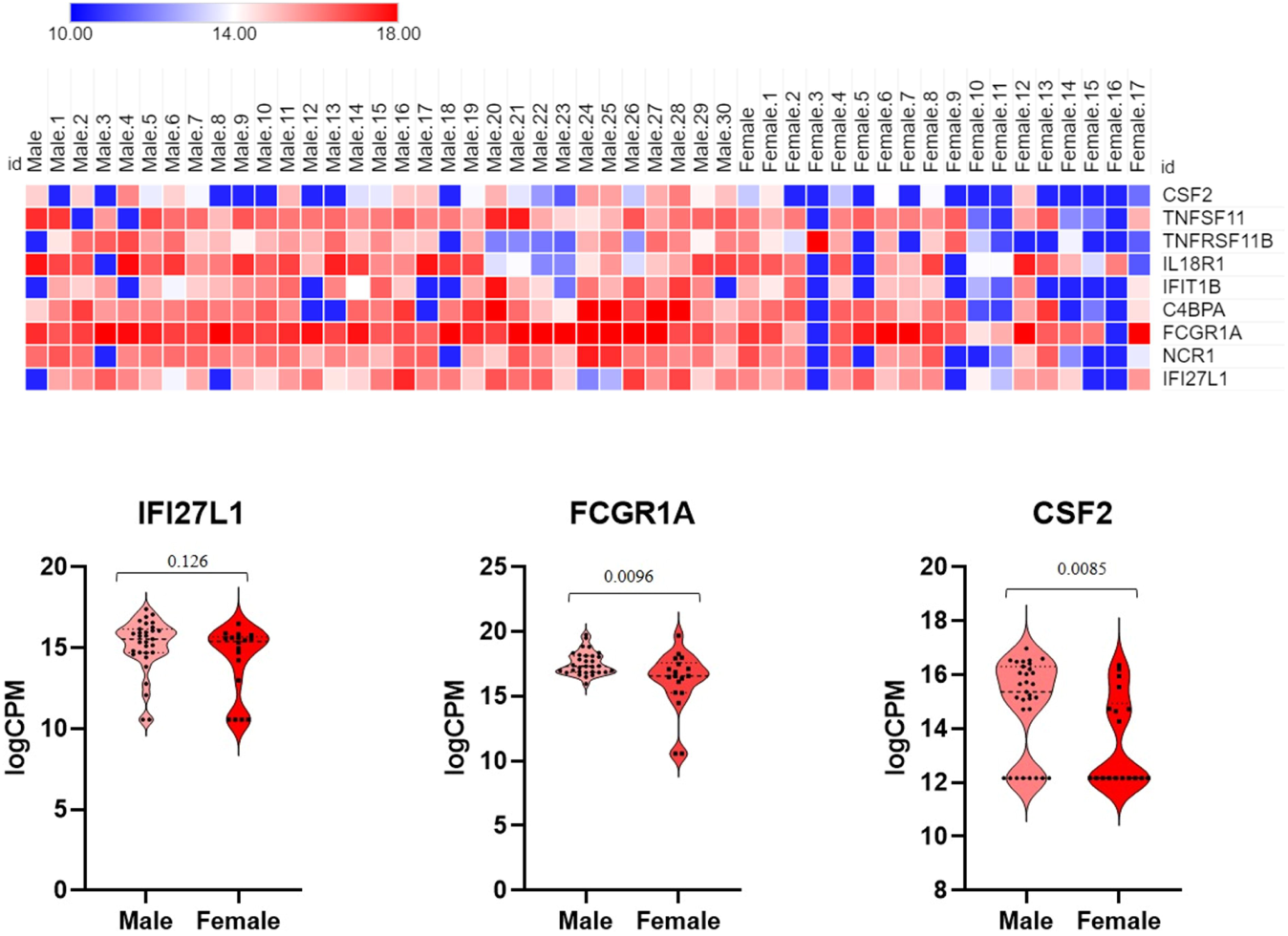
Dotplot visualization of upregulated (a), and downregulated pathways (b) in patients with mild, moderate and severe disease. The color of the dots represents adj *p*-value for each enriched pathways, and size represents the percentage of genes enriched in the total gene set. Volcano plots for upregulated and downregulated genes in patient with mild (c), moderate (d), and severe (e) outcomes. X-axis represents log_2_ fold change of genes and y axis represents –log_10_ *P*-value in DEG analysis.

