## Supplementary figures and images for "Host transcriptomic profiling of COVID-19 patients with mild, moderate, and severe clinical outcomes"

### Supplementary Figure 1

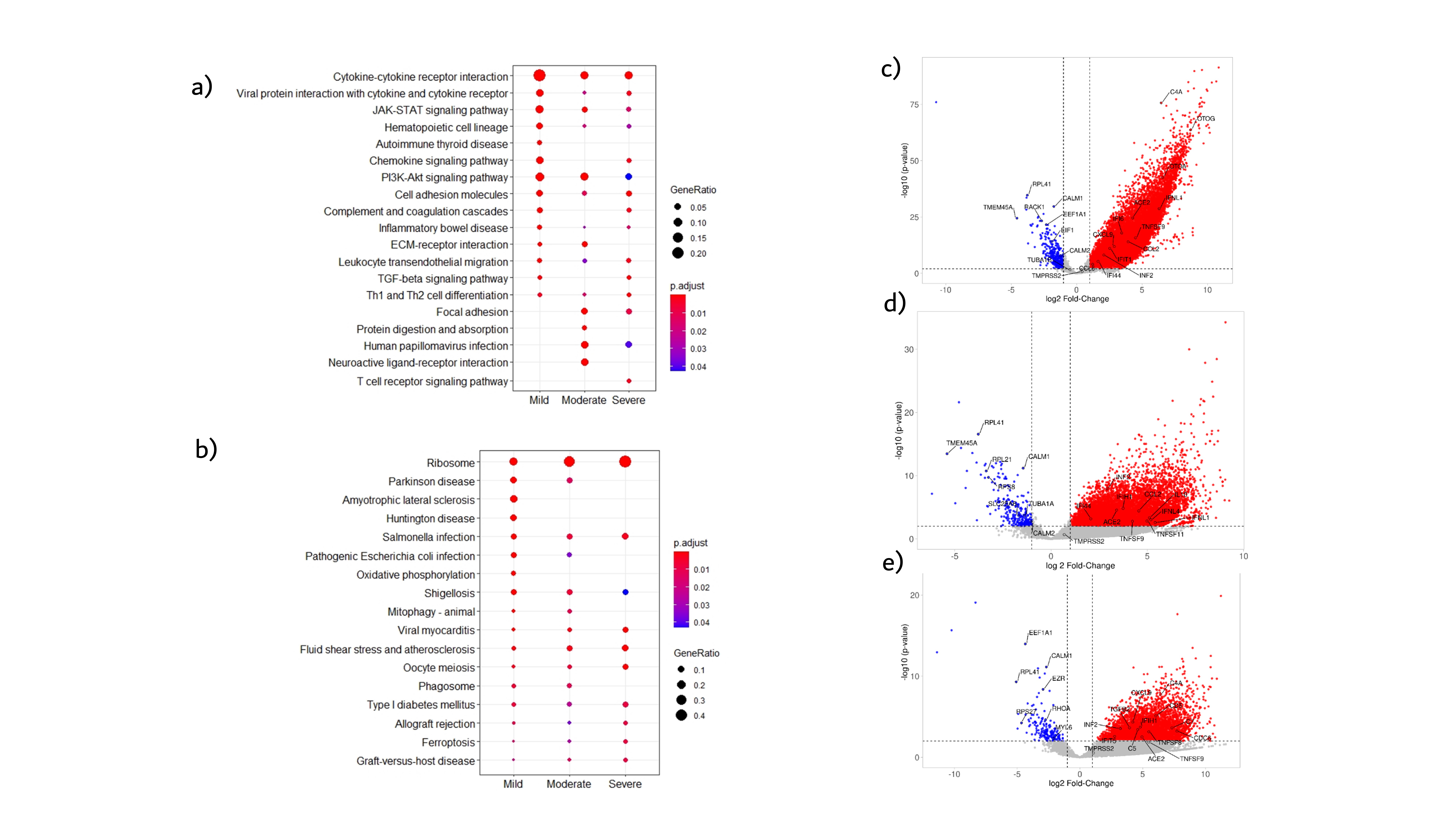
